# MaxGeomHash: An Algorithm for Variable-Size Random Sampling of Distinct Elements

**DOI:** 10.1101/2025.11.11.687920

**Authors:** Mahmudur Rahman Hera, David Koslicki, Conrado Martínez

## Abstract

With the surge in sequencing data generated from an ever-expanding range of biological studies, designing scalable computational techniques has become essential. One effective strategy to enable large-scale computation is to split long DNA or protein sequences into *k*-mers, and summarize large *k*-mer sets into compact random samples (a.k.a. *sketches*). These random samples allow for rapid estimation of similarity metrics such as Jaccard or cosine, and thus facilitate scalable computations such as fast similarity search, classification, and clustering. Popular sketching tools in bioinformatics include Mash and sourmash. Mash uses the MinHash algorithm to generate fixed-size sketches; while sourmash employs FracMinHash, which produces sketches whose size scales linearly with the total number of *k*-mers.

Here, we introduce a novel sketching algorithm, MaxGeomHash, which for a specified integer parameter *b* ≥ 1, will produce, without prior knowledge of *n* (the number of *k*-mers) a random sample of size *b* lg(*n/b*) + 𝒪(*b*). Notably, this is the first permutation-invariant and parallelizable sketching algorithm to date that can produce sub-linear sketches, to the best of our knowledge. We also introduce a variant, *α*-MaxGeomHash, that produces sketches of size *Θ*(*n*^*α*^) for a given *α* ∈ (0, 1). We study the algorithm’s properties, analyze generated sample sizes, verify theoretical results empirically, provide a fast implementation, and investigate similarity estimate quality. With intermediate-sized samples between constant (MinHash) and linear (FracMinHash), MaxGeomHash balances efficiency (smaller samples need less storage and processing) with accuracy (larger samples yield better estimates). On genomic datasets, we demonstrate that MaxGeomHash sketches can be used to compute a similarity tree (proxy for a phylogenetic tree) more accurately than MinHash, and more efficiently than FracMinHash. Our C++ implementation is available at: github.com/mahmudhera/kmer-sketch. Code to reproduce the analyses and experiments is at: github.com/KoslickiLab/MaxGeomHash.

## 1 Introduction

The exponential growth of genomic and metagenomic sequencing data has necessitated the development of scalable computational methods for efficiently processing and comparing biological sequences. Central to many methods is the concept of *k*-mers (substrings of length *k*), which serve as fundamental units for sequence comparison and analysis. However, the vast number of distinct *k*-mers in modern datasets makes exact comparisons computationally prohibitive, driving the need for approximation techniques that trade accuracy for improvements in speed and memory.

Sketching methods have emerged as a powerful solution, creating compact “fingerprints” of sequence data that preserve essential similarity information while reducing computational requirements. MinHash [1, 22] has become ubiquitous in genomic applications, enabling rapid all-versus-all comparisons through fixed-size sketches. However, as we and others have shown [16,17], traditional MinHash exhibits significant limitations when comparing sets of very different sizes, a common scenario in metagenomic analyses where bacterial genomes are compared against complex environmental metagenomic samples.

FracMinHash addresses these limitations by allowing sketch sizes to scale with the data [11, 13]. Rather than maintaining a fixed number of hash values, FracMinHash retains a fraction *s* ∈ [0, 1] of all *k*-mers. This linear scaling of sketch size enables accurate containment estimation between sets of arbitrary sizes.

While FracMinHash provides excellent accuracy, the price it pays is maintaining very large samples (linear in the number of distinct elements). In modern bioinformatics, genomic repositories and metagenomic studies often contain billions to trillions of sequences, resulting in very large sketch sizes. This motivates the need for new sampling algorithms that can achieve a middle ground: maintaining the accuracy benefits of letting samples grow, while keeping sample sizes sub-linear.

In this work, we propose a novel algorithm for random sampling, which we call MaxGeomHash (MGH for short), formulated as subsampling from a large finite dataset or data stream. Given a user-defined parameter *b*, our algorithm produces random samples of (expected) size *b* lg(*n/b*) + 𝒪 (*b*) from a data stream containing *n* distinct elements (where *n* is possibly unknown), providing a compelling balance between sample size and accuracy. We also present a variant, *α*-MaxGeomHash (*α*-MGH for short) that renders samples of expected size *Θ*(*n*^*α*^), *α* ∈ (0, 1), where *α* is a user-specified parameter. The parameter *b* in MaxGeomHash allows the user to control the sketch size while keeping the growth of sketches logarithmic, whereas the parameter *α* in *α*-MaxGeomHash allows for changing the asymptotic growth order of the sketch itself.

Importantly, MaxGeomHash and *α*-MaxGeomHash are the first dependable, order-independent, parallelizable, sub-linear sampling methods to date. Unlike existing sub-linear methods such as Affirmative Sampling [18], which is sensitive to the order of data stream processing and cannot be reliably parallelized, our approach produces identical samples regardless of data partitioning or thread execution order.

We theoretically characterize the expectation and variance of MGH and *α*-MGH sample sizes and quantify their computational cost rigorously. Our analyses also prove that MGH and *α*-MGH samples allow for unbiased or asymptotically unbiased similarity estimation, including Jaccard, cosine, and other metrics, by leveraging results from [19, 25].

Empirical evaluations on simulated and real genomic data confirm our theoretical predictions and demonstrate practical utility, including efficient pairwise similarity computation. We show that MGH and *α*-MGH allow for a balance between MinHash and FracMinHash, with provisions for a better accuracy-vs-efficiency tradeoff. Lastly, we provide an efficient C++ implementation for computing and comparing MGH/*α*-MGH samples directly from FASTA/FASTQ files, making our methods accessible to the bioinformatics community.

## 2 Background: Random Sampling from Data Streams

Random sampling has a long history in algorithms and data systems, from early treatments and classic textbook expositions [7, 14] to scalable streaming variants such as Reservoir Sampling [31] and its weighted forms [5]. In large-scale bioinformatics, sketching-based sampling is indispensable for scalable and efficient analysis. Prominent families include MinHash and bottom-*k* [1, 3], FracMinHash [9, 12], and Affirmative Sampling [18]; in biological contexts, FracMinHash-style sketches are particularly effective and can be used to estimate a variety of biologically relevant quantities [8, 9].

### Model and objective

We observe a long dataset or data stream *Ƶ* = *z*_1_, *z*_2_, …, *z*_*N*_ of items drawn from large *universe* 𝒟, with *z*_*i*_ ∈ 𝒟. Let the multiset of distinct elements underlying *Ƶ* be 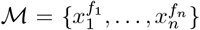, with *f*_*i*_ the frequency of *x*_*i*_ in *Ƶ*, and let the set of distinct elements be 𝒳 = {*x*_1_, …, *x*_*n*_}. The (unknown) number *n* = |𝒳 | is the *cardinality* of *Ƶ*. Our goal is to obtain a *random sample* 𝒮 of distinct elements from 𝒳 such that, for *S* = |𝒮|, every *S*-subset (a subset containing *S* elements) of 𝒳 is equally likely to be drawn. We focus on methods that operate in one pass over *Ƶ* without prior knowledge of *n*.

### A unifying hashed-sampling view

Most random sampling algorithms work by sequentially processing the dataset or data stream *Ƶ*, one element at a time, and keeping a current sample 𝒮 (in all the cases that we study here 𝒮 is a random sample from the subset of elements processed so far). If the current item *z* is present in 𝒮, we can update its frequency count and move on to the next. If *z* is not present, we compute its hash value *h* = *h*(*z*) and use *h* to decide whether *z* is added to the sample or discarded. Moreover, and depending on our decision w.r.t. *z*, we might decide to make additional changes in 𝒮, for example, removing some elements from 𝒮. Examples of algorithms that fall into the general scheme outlined above include bottom-*k* and MinHash [1]^5^ FracMinHash [8, 9, 12], ModHash [1], and Affirmative Sampling [18].

In FracMinHash, an item *z* is sampled (kept in the sample 𝒮) iff *h*(*z*) ≤ *s* for a given real value *s* ∈ (0, 1), where *h*(.) produces real hash values in the range [0,1]. In ModHash, an item *z* is sampled iff *h*(*z*) mod *M* = 0, where *h*(.) produces integer hash values in some range [0, *H*] for some large *H*. In bottom-*k*, an item *z* is sampled iff it is among the *k* smallest hash values seen so far. If sampling *z* results in |𝒮|= *k* + 1, then the item with the largest hash value in 𝒮 is removed. MinHash applies *k* independent hash functions, and an item *z* is sampled if *h*_*i*_(*z*) is the minimum hash value among the *h*_*i*_ values observed so far, for some *i*. Affirmative Sampling —the standard variant— samples an item *z* if not present in the sample and if *h*(*z*) is larger than min_*x*∈𝒮_ *h*(*x*); moreover, it evicts the item *z*^∗^ with the smallest hash value in 𝒮 if *h*(*z*) is not among the *b*-largest hash values in 𝒮. Initially, the first *b* items are always included in 𝒮; and at any given moment, the sample 𝒮 will contain |𝒮| distinct elements with largest hash values seen so far.

In FracMinHash and ModHash, the expected size of the sample 𝒮 is linear in *n*. For FracMinHash, it will follow a binomial distribution of parameters *n* and *s*. Likewise, ModHash produces random samples with their size following a binomial distribution of parameters *n* and 1*/M*. Last but not least, in Affirmative Sampling, the expected size of 𝒮 is ∼ *b* ln(*n/b*), for a fixed parameter *b*, in the standard variant; in *α*-Affirmative Sampling, 0 *< α <* 1, the expected size of the sample is *Θ*(*n*^*α*^).

MinHash has the advantage of producing samples of fixed size and the samples/sketches are small and easy to obtain. However, inference based on those sketches is less accurate, especially when they are used to estimate containment and similarity indices between two sets. Conversely, FracMinHash and ModHash give excellent accuracy, but the price they pay is to keep very large samples (linear in the number of distinct elements). Midway, we have Affirmative Sampling, which guarantees very good accuracy and (asymptotically) unbiased estimation, while producing significantly smaller samples (of size *Θ*(log *n*) or 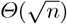, say).

All these algorithms return random samples of distinct elements, which is easily provable under pragmatic assumptions on the randomness and independence of the hash functions. Indeed, if all decisions (to sample/to evict/to discard) are solely based upon the hash values of the items, then the samples will be random.

### Dependability

A sampling algorithm is *dependable* [18] if it allows for exact frequency counts: membership of *x*_*j*_ in 𝒮 is fixed at its first occurrence, and items once removed are never reinserted. Reservoir sampling [31] is not dependable, whereas all the other algorithms mentioned are.

### Parallel composition

Massive datasets must be processed in parallel by partitioning *Ƶ* into *p* disjoint data subsets or substreams *Ƶ*_1_, …, *Ƶ*_*p*_. For order-independent sketches there is a standard merge operation: compute a local sample 𝒮_*i*_ from each *Ƶ*_*i*_ and then aggregate 𝒮 = *ϕ* (∪_1≤*i*≤*p*_ 𝒮_*i*_, where *ϕ* is the algorithm-specific pruning rule (e.g., keep the *k* smallest hash values, or keep all items with *h*(*x*) ≤ *s*). This equivalence works both ways: running the algorithm on the entire stream *Ƶ* or merging local sketches yields the same distribution. MinHash, bottom-*k*, FracMinHash, and ModHash satisfy this property; Reservoir Sampling (with repetitions) and Affirmative Sampling (both variants) do not, and are not mergeable in this sense.

### Expected cost of random sampling

The *cost* of running a sampling algorithm refers to the number of operations required to scan the entire data stream and produce the random sample. We give the expected cost *M* (*n, N*) of the random sampling algorithms in the form *Θ*(*N* + *u*(*n*) · *t*(*S*)), where *N* is the size of the dataset (or the length of the data stream), *n* the number of distinct elements (*n* ≤ *N*, in many applications *n* ≪ *N*), *u*(*n*) is the expected number of times there is an update of the sample (and of auxiliary data structures, if any), and *t*(*S*) is the time to update a sample of size *S*. This cost is based on the assumption that we can test whether a given item *z* (or hash value *h* = *h*(*z*)) is already present in the sample or not in expected constant time. The expected number of updates *u*(*n*) will be at least *S*, but it is often the case that *u*(*n*) ≫ *S*, for example, in bottom-*k* we have *S* = *k*, but *u*(*n*) = *k* ln(*n/k*) + 𝒪 (*k*) [18]; using a min-heap to keep and efficiently update the sample we have *t*(*S*) = 𝒪 (log *k*). This gives a total expected cost for bottom-*k* of 𝒪 (*N* + *k* log *k* log(*n/k*)). Alternately, the cost of producing a FracMinHash sketch is 𝒪 (*N* + *ns*), as *u*(*n*) = *S* = *n* ·*s* and *t*(*S*) = *Θ*(1). The expected cost of the new algorithms, MGH and *α*-MGH, will be investigated in Sections 3 and 4, respectively, where we describe and study their main properties.

### At-a-glance comparison

We summarize the main properties that guide the choice of a sampler in practice. Dependability is necessary if exact frequency counts are required; mergeability enables distributed processing; the parameter controls the memory footprint; and the last two columns report the expected sample size. Asterisks indicate quantities that are either fixed or bounded by a constant, and *n* denotes the total number of distinct elements in the original set.

| Algorithm | Dependable? | Mergeable? | Parameter | Expected $\vert \mathcal{S}\vert $<br>(parameterized) | Expected $\vert \mathcal{S}\vert $<br>(fixed parameter) |
| --- | --- | --- | --- | --- | --- |
| MinHash [1] | ✓ | ✓ | $k \in \mathbb{N}$ | $k$ (*) | $\mathcal{O}(1)$ |
| ModHash [1] | ✓ | ✓ | $m \in \mathbb{N}$ | $n/m$ | $\mathcal{O}(n)$ |
| Reservoir Sampling<sup>6</sup> [31] | ✗ | ✗ | $k \in \mathbb{N}$ | $\leq k$ (*) | $\mathcal{O}(1)$ |
| FracMinHash [12] | ✓ | ✓ | $s \in (0, 1)$ | $n \cdot s$ | $\mathcal{O}(n)$ |
| Affirmative Sampling [18] | ✓ | ✗ | $b \in \mathbb{N}$ | $b \ln(n/b) + \mathcal{O}(b)$ | $\mathcal{O}(\log n)$ |
| α-AS [18] | ✓ | ✗ | $\alpha \in (0, 1)$ | $\Theta(n^\alpha)$ | $\mathcal{O}(n^\alpha)$ |
| MGH (this paper) | ✓ | ✓ | $b \in \mathbb{N}$ | $b \log_2(n/b) + \mathcal{O}(b)$ | $\mathcal{O}(\log n)$ |
| α-MGH (this paper) | ✓ | ✓ | $\alpha \in (0, 1)$ | $\frac{2^{1/(1-\alpha)}-1}{2^{\alpha/(1-\alpha)}-1} \cdot n^\alpha + \mathcal{O}(n^\alpha)$ | $\mathcal{O}(n^\alpha)$ |

In summary, our algorithm fits neatly between the fixed-length samples of MinHash and the linearly-sized samples of FracMinHash, while retaining mergeability (which is absent from Affirmative Sampling).

## 3 MaxGeomHash

Here, we describe our algorithm, MaxGeomHash (MGH for short) which is dependable and parallelizable in the sense explained in the previous section. Like Affirmative Sampling, it has the nice property of returning samples of variable size, growing with *n*. In particular, the standard variant of the algorithm, which we examine now, returns samples of expected size *b* lg(*n/b*) + 𝒪 (*b*)—here, and in the sequel we use lg ≡ log_2_.

Given a bit string *w* ≠ 000 · · · 0, let zpl(*w*) denote the length of the longest prefix of 0s in *w* (if *w* = 0000 · · · then zpl(*w*) = | *w* |, by convention). In other words, if zpl(*w*) = *j* then the first *j* bits of *w* are 0 (bits 0 to *j* − 1), and bit *j* is 1. We will also use tail(*w, j*) to denote the bit string that results from removing the first *j* + 1 bits of *w* (that is, the longest prefix of zeros and the leftmost 1).

For each data stream item *z*, we compute its hash value *h* := *h*(*z*), and determine the position *i* = 1+zpl(*h*) of the leftmost 1 in the binary representation of *h*. Here we assume that the hash function *h*(·) maps an item *z* to a *ℓ*-bit positive integer *h*(*z*) in the range [0, *H* − 1] for some large *H* = 2^*ℓ*^ (in practice, we used *ℓ* = 64). We also set *h*^′^ as the suffix after the leftmost 1. We use *i* to index *z* into a bucket 𝒮_*i*_. If 𝒮_*i*_ already contains *z* or *h*^′^ is not among the *b* largest hash values in 𝒮_*i*_ then we will discard *z*. If 𝒮_*i*_ doesn’t yet contain *b* elements or *h*^′^ is among the *b* largest hashes in 𝒮_*i*_ then we add *z* to 𝒮_*i*_; in the latter case, the element *z*^∗^ with the smallest hash value *h*^′^ is evicted, so that 𝒮_*i*_ will contain the ≤ *b* elements in the data stream such that their leftmost 1 is at position *i* of *h*(*z*) and have largest *h*^′^ = tail(*h*(*z*), *i*). A frequency counter is kept for all *z* ∈ 𝒮. The steps of MaxGeomHash are detailed in Algorithm 1. A visual overview of MaxGeomHash is given in Figure 1.

**Fig. 1:**
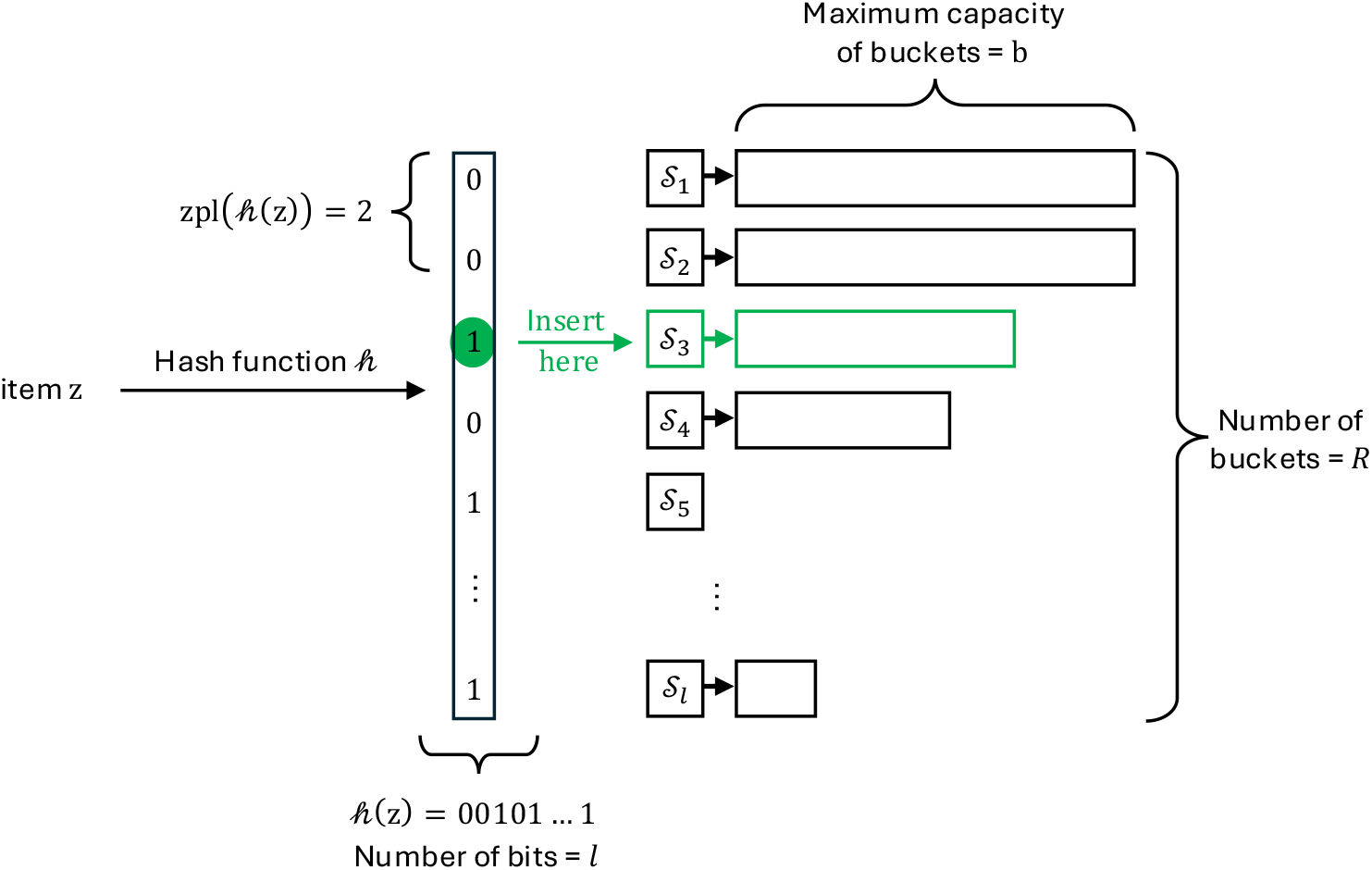
Overview of MaxGeomHash. A hash function *h*(·) maps an item *z* to a bit string of length *ℓ*. The zero prefix length (zpl) of the hash value *h*(*z*) determines in which bucket *i* we should sample (or not) *z*. Each bucket 𝒮_*i*_ ≡ bucket[*i*] has a maximum capacity of *b*, which is the integer parameter of MaxGeomHash, where *b* ≥ 1. Of all items assigned to a bucket, only the items with the *b* largest hash values are retained (similar to bottom-*k* sketches, only we are keeping the largest hash valued items. The number of buckets can be at most *ℓ*, the number of bits in the hash value, but this will never happen in practice if lg *n* ≪ *ℓ*. The largest index *R* of a non-empty bucket is given by the maximum of *n* i.i.d. geometric random variables, hence the “MaxGeom” in the algorithm’s name.

### Algorithm 1

MaxGeomHash algorithm

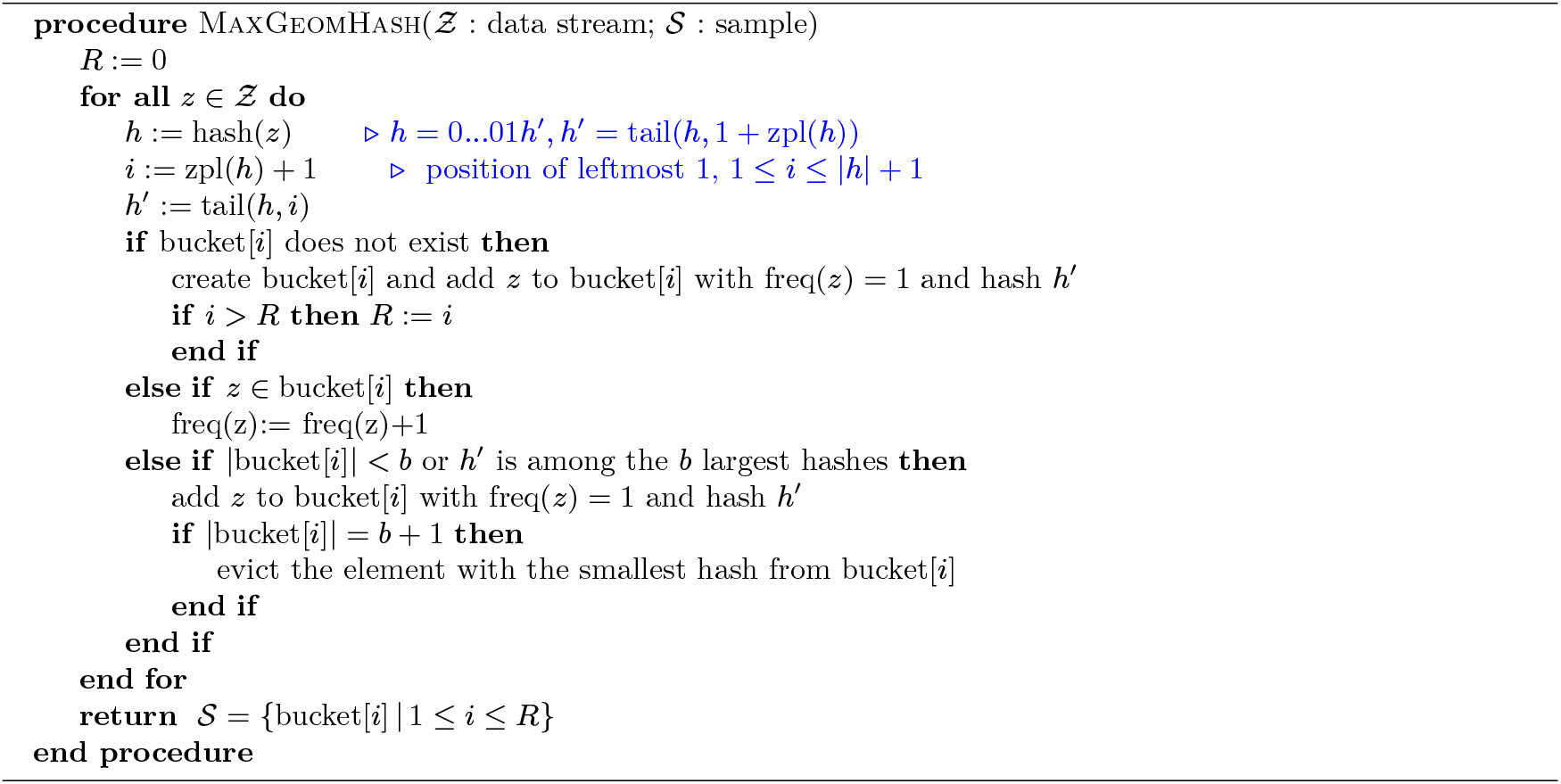

Let *x*_1_, …, *x*_*n*_ denote the distinct elements in *Ƶ*. Let *ρ*_*i*_ = 1 + zpl(hash(*x*_*i*_)). The *ρ*_*i*_ are i.i.d. geometric random variables with parameter 1/2. Therefore, the probability that an item *z* has to be considered for bucket 𝒮_*i*_ is 1*/*2^*i*^. Using this, we can compute the expected sample size and variance.

### Theorem 1.

*M**ax**G**eom**H**ash* *with parameter b* ≥ 1 *will produce a random sample* 𝒮 *of size S* = |𝒮| = ∑_1≤*i*≤*R*_ |𝒮_*i*_| *such that*

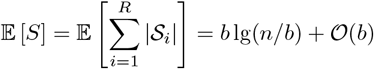

*where R* = max{*ρ*_*j*_ | 1 ≤ *j* ≤ *n*}. *Moreover*, 𝕍 [*S*] = *Θ*(1).

*Proof (Sketch of the proof)*. We give here only an intuition of the proof for 𝔼 [*S*]. A detailed proof will be available in the full version of this paper.

Each bucket 𝒮_*i*_ receives (but only keeps up to *b*) a certain number *X*_*i*_ ∼ Binomial(*n*, 1*/*2^*i*^) of items from the data stream *Ƶ*, that is, *S*_*i*_ = |𝒮_*i*_| ∼ min(*b*, Binomial(*n*, 1*/*2^*i*^)) and the indices of buckets run from 1 to *R*, with *R* = max{*ρ*_1_, …, *ρ*_*n*_}. We know 𝔼 [*R*] = lg *n* + 𝒪(1) and 𝕍 [*R*] = *Θ*(1) [29]. For the size of the final sample 𝒮 we have 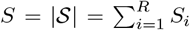, and because of linearity of expectations and the fast decay of the terms with *i* ≥ lg *n*, it is not difficult to prove that

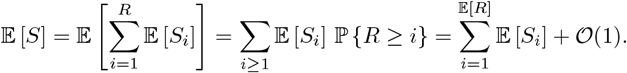

For *i* ≲ *i*^∗^ := lg(*n/b*) we will have *S*_*i*_ = *b* with high probability; likewise, for *i* ≳ *i*^∗^, *S*_*i*_ = min{*X*_*i*_, *b*} = *X*_*i*_ with high probability, hence 𝔼 [*S*_*i*_] = *n/*2^*i*^ and so 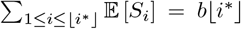, and 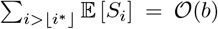. The terms around the critical index *i*^∗^ and the error made by assumming min(*X*_*i*_, *b*) = *b* for *i* ≤ ⌊*i*^∗^⌋ and min(*X*_*i*_, *b*) = *X*_*i*_ for *i >* ⌊*i*^∗^⌋ are also 𝒪 (*b*) (actually, *o*(*b*)).

For the variance, we use a similar approach, considering three regions. We notice that Cov [*S*_*i*_, *S*_*j*_] = 0 with high probability if *i* or *j* are below *i*^∗^, they are exponentially small if *i* ≫ *i*^∗^ and *j* ≫ *i*^∗^ (then *X*_*i*_ and *X*_*j*_ behave as independent random variables and thus Cov [*S*_*i*_, *S*_*j*_] ≈ *n/*2^*i*+*j*^), and only a constant number of terms Cov [*S*_*i*_, *S*_*j*_] with *i* ≈ *i*^∗^ and *j* ≈ *i*^∗^ contribute to 𝕍 [*S*].

The expected cost *M*_*b*_(*n, N*) of MGH when processing a data stream of *N* items such that *n* of them are distinct is dominated by the time needed to scan the data stream 𝒪 (*N*), plus the time to update the buckets *i* that contain *b* elements. We expect *N/*2^*i*^ items to be directed to bucket *i*, but most will be discarded with cost *Θ*(1), as they are repetitions or because their hash *h*^′^ is not among the *b* larger in 𝒮_*i*_. Like in a bottom-*k* approach, we expect ∼ *b* ln(*n/b*2^*i*^) updates in bucket 𝒮_*i*_, and the cost of each of these updates is 𝒪 (log *b*). Summing over all possible *i*, we get that the expected cost is

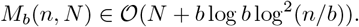

## 4 *α*-MaxGeomHash

We now discuss a variant of MGH, called *α*-MGH, that renders samples of expected size *Θ*(*n*^*α*^) for a fixed parameter *α* ∈ (0, 1) of our choice. The algorithm is exactly as Algorithm 1, but instead of keeping at most *b* distinct elements on each 𝒮_*i*_, we will keep up to ⌈2^*βi*^⌉ in bucket *i*, with 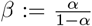. Additional fine tuning can be introduced by setting the maximum capacity of each bucket to *γ* · 2^*βi*^, for a given constant *γ >* 0, but for the rest of the section we will stick to *γ* = 1.

The intuition is that bucket *i*, for small *i*, namely, *i < i*^∗^ = (1 − *α*) lg *n*, will receive a lot of elements (∼ *n/*2^*i*^) but only will keep very few (2^*βi*^ ≪ *n/*2^*i*^), whereas for large *i, i > i*^∗^, there are very few items going to bucket *i* but all of them will be kept (*n/*2^*i*^ ≪ 2^*βi*^). For *i*^∗^ = (1 − *α*) lg *n*, and since *α* = *β/*(1 + *β*) and 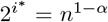, it follows that 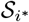 will contain at most 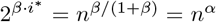 elements, but on the other hand we expect that *n/n*^1−*α*^ = *n*^*α*^ distinct elements are directed to bucket *i*^∗^ and kept since 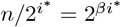. Hence 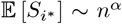. The accumulated contribution of the other buckets with values of *i* ≠ *i*^∗^ can be shown to be 𝒪(*n*^*α*^). Actually, a more delicate analysis yields the following theorem.

### Theorem 2.

*The variant α-MGH of M**ax**G**eom**H**ash* *with parameter α* ∈ (0, 1) *will produce a random sample* 𝒮 *of size S such that*

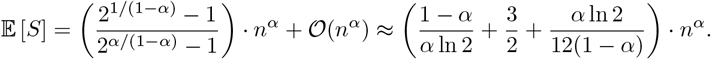

*Moreover* 𝕍 [*S*] = *Θ*(*n*^*α*^).

The proof will be available in the full version of this preprint and runs along similar lines as that of Theorem 1. The 𝒪 (*n*^*α*^) accounts for a small adjustment needed when { (1 − *α*) lg *n* }≠ 0 (we use {*x*} = *x*−⌊*x*⌋ to denote the fractional part of *x*).

We can also express expected cost *M*_*α*_(*n, N*) in the same form as we have discussed for the other algorithms, namely, *M*_*α*_(*n, N*) = 𝒪 (*N* + *u*(*n*)*t*(*S*)). Since the expected total number of updates *u*(*n*) is 𝒪 (*n*^*α*^) and we can bound the cost of updating any bucket by 𝒪 (log(*n*^*α*^)) = 𝒪 (log *n*), one therefore has the expected cost of

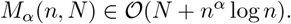

## 5 Similarity estimation with MGH and *α*-MGH

Given two sets *A* and *B* and random samples 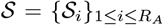 and 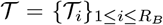, we will use them to estimate Jaccard(*A, B*) = | *A* ∩ *B* | */* | *A* ∪ *B* |. We use *A*_*i*_ and *B*_*i*_ to denote the subsets of *A* and *B* of elements *x* with zpl(*x*) + 1 = *i*. If we use the same hash function to sample from *A* and from *B*, then if an item *x* ∈ *A* is considered for bucket 𝒮_*i*_ then it will be also considered for bucket 𝒯_*i*_ if also appears in *B*, and vice versa. By definition, we have *A*_*i*_ ∩ *A*_*j*_ = ∅ whenever *i* ≠ *j*, and U _*i* ≤1_ *A*_*i*_ = *A*; likewise,{ *B*_*i*_ } _*i*≥1_ gives a partition of *B*. One important result [19] establishes that if *A* ∩ *B* = ∅ then 𝒮+ 𝒯 is a random sample of *A* + *B* (this also holds for 𝒮_*i*_ + 𝒯_*i*_). But 𝒮_*i*_ ⋃ 𝒯_*i*_ is not a random sample of *A*_*i*_ ∪ *B*_*i*_, nor𝒮 ∪ 𝒯 a random sample of *A* ∪ *B*. Therefore, we need to apply a “filtering” or “refining” step. In particular, we will keep only the *b* elements with larger hash values (à la bottom-*k*). So for each *i*, we will compute 𝒮_*i*_ ∪ 𝒯_*i*_ but keep the *b* elements with larger hash values *h*^′^ (or only |𝒮_*i*_ ∪ 𝒯_*i*_| elements if that cardinality is smaller than *b*). Let us call this new sample 𝒰_*i*_. This is indeed a random sample of *A*_*i*_ ∪ *B*_*i*_. Moreover, we will compute 𝒱_*i*_ = 𝒮_*i*_ ∩ 𝒯_*i*_. Notice that |𝒱_*i*_| ≤ *b*, since both 𝒮_*i*_ and 𝒯_*i*_ contain at most *b* elements each.

All 𝒰_*i*_ are mutually disjoint (𝒰_*i*_ ∩ 𝒰_*j*_ = ∅ if *i* ≠ *j*) which entails that 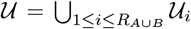 is a random sample of *A* ∪ *B*. Likewise 𝒱 = U_*i*_ 𝒱_*i*_ = *A* ∩ *B* ∩ 𝒰, hence (see [19] for a detailed proof) |𝒱|*/*|𝒰 | is an unbiased estimator of Jaccard(*A, B*). In fact, we can use 𝒰 and 𝒱 to estimate many other set similarity and set containment measures [19].

The arguments also work *verbatim* if instead of buckets with maximum capacity equal to *b*, their capacity is given by *f*_*i*_, e.g., *f*_*i*_ = ⌈2^*βi*^ ⌉, as in *α*-MGH. When computing 𝒰_*i*_ we would just need to do the unions 𝒮_*i*_ ∪ 𝒯 _*i*_, then retain the elements with the *f*_*i*_ larger *h*^′^ values (or less if there are not *f*_*i*_ elements in 𝒮_*i*_ ∪ 𝒯 _*i*_).

In order to merge two samples 𝒮 and 𝒯 we should proceed as described above, taking advantage that the two samples are already classified into buckets, and computing the unions/intersections of the buckets 𝒮_*i*_ and 𝒯_*i*_ while only keeping the *b* (or *f*_*i*_ = ⌈ 2^*βi*^⌉) elements with larger *h*^′^ hash values, doing it in one single pass. But from the point of view of the result, the process is equivalent to applying MGH (or *α*-MGH) to 𝒮 ∪ 𝒯, or 𝒮 ∩ 𝒯, to get the union or the intersection of the two samples; indeed, we have the following important property^7^:

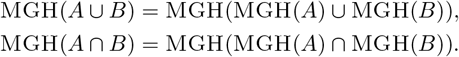

Notice that MinHash and FracMinHash satisfy the analogous equalities.

From there, the results of [9, 19, 25] apply, showing that we can use MGH or *α*-MGH samples to get (asymptotically) unbiased estimates of the Jaccard similarity of *A* and *B*, of their cosine similarity, of the containment index, and many other similarity measures, like Kulczynski 1, Kulczynski 2, or Sørensen-Dice. A list and definition of these metrics can be found here [19, Table 3].

For some similarity measures like Jaccard, containment index, or Kulczynski 2, the estimates are unbiased; for others, like cosine or Kulczynski 1, the bias is 𝒪 (𝔼 [1*/S*]), with *S* the size of the sample—which is a random variable for MGH and *α*-MGH (and also for FracMinHash). Also, the variance for all the different similarity measures mentioned above is 𝒪(𝔼 [1*/S*]).

### Theorem 3.

*Let σ*_*u*_ *be any of the following similarity measures: Jaccard, containment, Kulczynski 2, Sørensen-Dice, or correlation coefficient. Then*

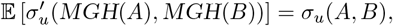

*where* 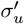 *is defined as σ with a final “filtering” step. For example, for Jaccard we have* 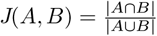 *and* 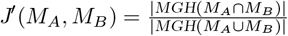. *The filtered versions of the other similarity measures are defined analogously. Notice that MGH*(*M*_*A*_ ∩ *M*_*B*_) = *M*_*A*_ ∩ *M*_*B*_, *and thus the filtering in the numerator can actually be avoided*.

*Let σ*_*b*_ *denote the cosine similarity, or Kulczynski 1. Then* 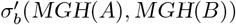 *is asymptotically unbiased:*

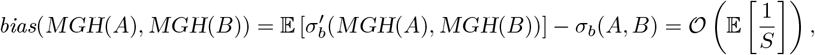

*with S* = |*MGH*(*MGH*(*A*)∪*MGH*(*B*))|. *Finally, for any similarity measure σ among those mentioned above*

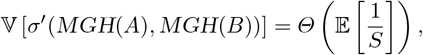

*where S is as above*.

Hence the bias, if any, the variance, and the mean square error (MSE=bias^2^+variance) of all our estimates based on MGH or *α*-MGH samples will all go to 0 because 𝔼 [1*/S*] = 1*/*𝔼 [*S*] + *o*(1*/*𝔼 [*S*]) and 𝔼 [*S*] → ∞ if *n* → ∞. Details of the proof that 𝔼 [1*/S*] ∼ 1*/*𝔼 [*S*] can be found in Appendix A. In general, it is not true, but because 𝔼 [*S*] → ∞, plus 𝕍 [*S*] = 𝒪 (𝔼 [*S*]), it is not difficult to obtain such remarkable result.

Irrespective of how we produce the random samples to be used in similarity estimation, the cost of computing such estimates will be linear in the sizes of the samples (plus the time needed to sort the samples, but this is only paid once, or can be omitted if we use hash-join to compute intersections and unions). Therefore, when we have smaller sketches (like in bottom-*k*) the computation of similarity estimates is extremely fast, but can be quite inaccurate. On the contrary, when the sketches are very large (like in FracMinHash) the estimates are very accurate, but the computation of the estimate is very time-consuming (see, for example, Table 1 in Section 6.5). MGH and *α*-MGH offer appealing compromises of good accuracy and moderate computational resources.

**Table 1:** Computational resources used by four sketching algorithms in computing the similarity tree of ten mammals. The trees are shown in Figure 6.

| Algorithm name | Computing sketches |  | Storing sketches | Computing all-pair Jaccard |  |
| --- | --- | --- | --- | --- | --- |
|  | CPU Time (s) | Peak Memory Usage (GB) | Disk space | CPU Time (s) | Memory (MB) |
| bottom-$k$ MinHash ($k = 1000$) | 4673.3 | 1.397 | 160 KB | 0.01 | 4.203 |
| FracMinHash ($s=0.001$) | 3970.84 | 1.421 | 360 MB | 20.64 | 977.621 |
| MAXGEOMHASH ($b=90$) | 4365.62 | 1.397 | 878 KB | 0.04 | 5.836 |
| α-MAXGEOMHASH ($\alpha=0.45$) | 4465.38 | 1.399 | 18 MB | 0.94 | 43.953 |

**Table 2:** Mammalian reference genomes downloaded from Ensembl (release 112) using wget.

| Standard name | Scientific name | Ensembl assembly ID |
| --- | --- | --- |
| Human | *Homo sapiens* | GRCh38 |
| Chimpanzee | *Pan troglodytes* | Pan_tro_3.0 |
| House mouse | *Mus musculus* | GRCm39 |
| Norway rat | *Rattus norvegicus* | mRatBN7.2 |
| Dog | *Canis lupus familiaris* | ROS_Cfam_1.0 |
| Cat | *Felis catus* | Felis_catus_9.0 |
| Cow | *Bos taurus* | ARS-UCD1.3 |
| Pig | *Sus scrofa* | Sscrofa11.1 |
| Horse | *Equus caballus* | EquCab3.0 |
| Gray short-tailed opossum | *Monodelphis domestica* | ASM229v1 |

## 6 Experiments and Results

In this section, we present: 1) simulation experiments verifying theoretical expectations and asymptotically unbiased estimators; 2) a real data experiment highlighting the balance provided by our algorithms.

### 6.1 MaxGeomHash samples grow sub-linearly with the original set

First, we validate the theoretical expectations of MGH and *α*-MGH sample sizes using simulated experiments. In these experiments, we generated random sets of 10K to 1M elements consisting of randomly generated alphanumeric strings of length 10. We generated MGH samples with parameters *b* ∈ { 70, 80, 90, 100 }, and *α*-MGH samples with parameters *α* ∈ { 0.4, 0.45, 0.5 }. For each choice of *b* and *α*, we ran 50 independent trials using different hash seeds and recorded the resulting sample size. The average sample sizes from these 50 trials are shown in Figure 2. The theoretical expectations outlined in Theorems 1 and 2 are also shown in Figure 2 using dashed black lines. When plotting these expected sizes, we excluded the smaller order terms (specifically, *ϵ*_*n,b*_ for the case of MGH, and 𝒪 (log *n*) for the case of *α*-MGH) from calculation.

**Fig. 2:**
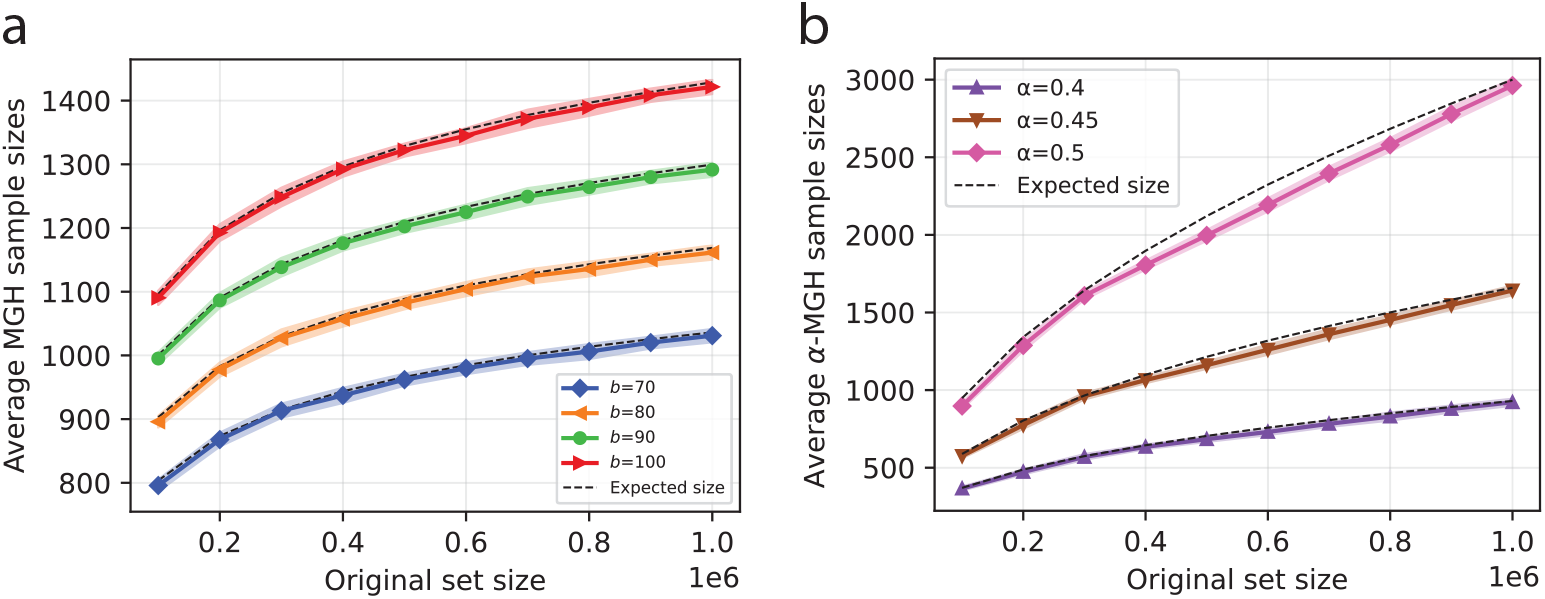
Average sample sizes, recorded from 50 independent trials. Samples are computed using MGH (averages shown in (a)) and *α*-MGH (averages shown in (b)). The solid lines show the averages, and the shaded regions around the solid lines (very narrow) highlight one standard deviation. Theoretical expectations (excluding lower-order terms) are shown in dashed black lines, closely tracked by the average recorded from simulations.

The plots clearly show that the growth of the MGH and *α*-MGH samples aligns closely with the theoretically derived expectation. As the size of the original set grows, the sample size scales sub-linearly, as we expect them to. Figure 2 also shows one standard deviation as shaded regions around the corresponding solid lines. Notably, the variance we observe for MGH and *α*-MGH samples is quite small – which highlights the stability of the size of the samples produced by MaxGeomHash.

### 6.2 MaxGeomHash provides more stability than Affirmative Sampling

As outlined in Section 2, the only other sub-linear (super-constant) sketching algorithm is Affirmative Sampling. In this section, we show that MGH sketches are more stable than sketches computed using Affirmative Sampling. We use AS to denote Affirmative Sampling henceforth. There are two versions of Affirmative Sampling – the plain version is referred to as AS, and the *α*-version (being able to produce sketches of size 𝒪 (*n*^*α*^), in expectation) is referred to as *α*-AS.

For the comparisons we present in this section, we generated a random set of 100,000 elements (each a random alphanumeric string) and computed sketches of it using AS and MGH, and the *α*-variant of both algorithms. Sketch sizes are shown in Figure 3a-d. We used *k* = 70 to run AS, *b* = 70 to run MGH, and used *α* = 0.4 to run both *α*-variants. These parameters were selected arbitrarily; any selection of other values would also reveal similar results.

**Fig. 3:**
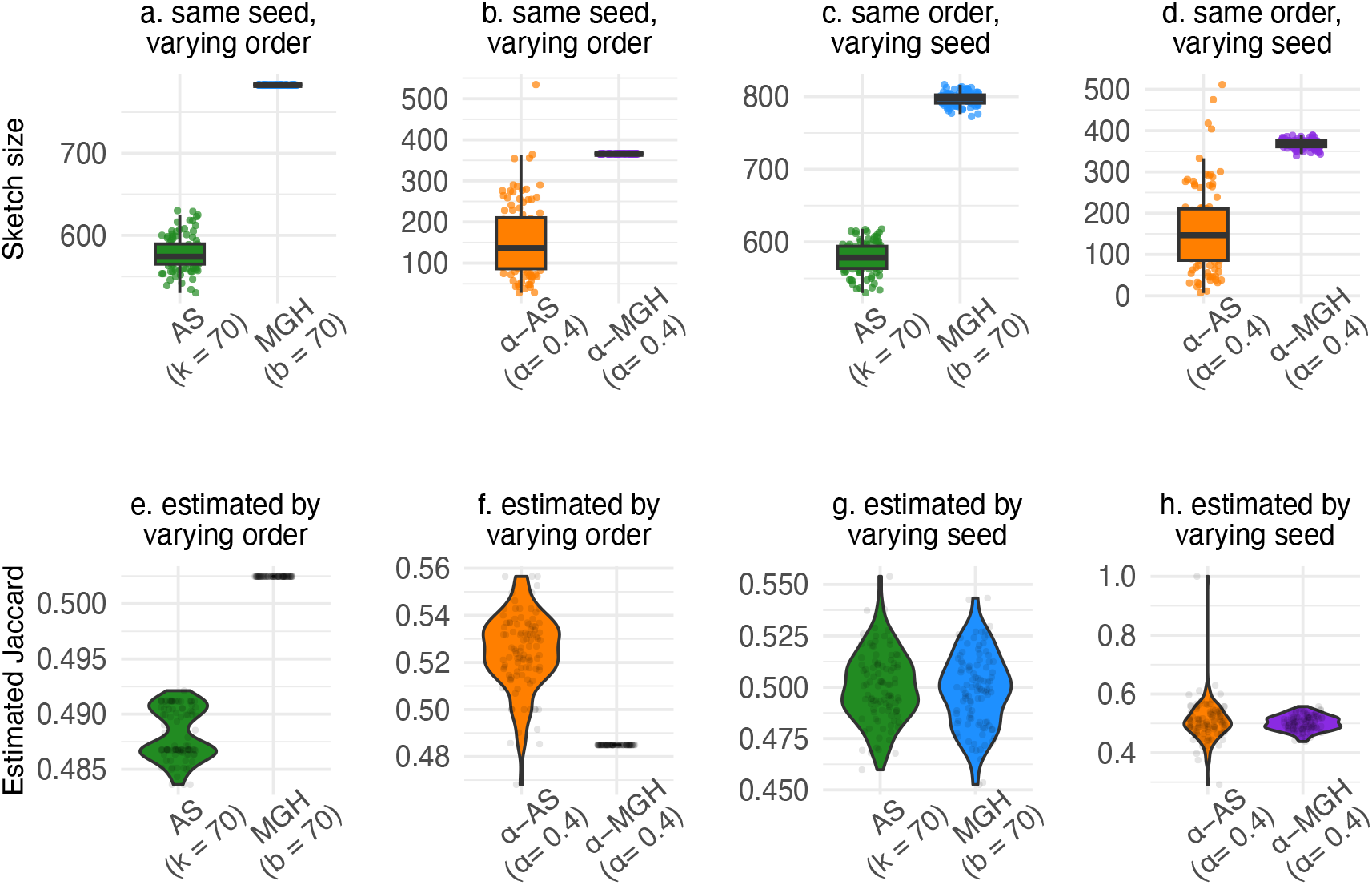
Comparison of AS and MGH sketches on random sets. (a–b) Sketch sizes across 100 random processing orders (fixed hash): AS and *α*-AS are order-sensitive, while MGH and *α*-MGH are order-invariant. (c–d) Sketch sizes across 100 hash functions (fixed order): MGH and *α*-MGH show substantially lower size variability than AS and *α*-AS, with *α*-AS being particularly unstable. (e–f) Jaccard estimates (true value 0.5) across random orders: MGH gives identical estimates, whereas AS (especially *α*-AS) varies widely. (g–h) Jaccard estimates (true value 0.5) across hash functions: AS and MGH show similar variability, but *α*-AS remains highly unstable.

First, when we ran the sketching algorithms, we processed the elements in 100 random orders. Figure 3a and Figure 3b show that simply by processing the elements in a different order, AS and *α*-AS produce different sketches despite using the same hash function (controlled by the random seed). On the other hand, MGH and *α*-MGH produce the same sketch each time. This “order invariant” property exhibited by MGH is important for reproducing the same results when rerunning experiments and plays a crucial role in multithreaded programming, where the order in which elements are processed often cannot be determined beforehand.

We next investigate the sketches generated by AS and MGH when the elements are always processed in the same order, but the sketches are generated with 100 different hash functions (each with a different seed). We show the sizes of AS and MGH sketches in Figure 3c, and the sizes of *α*-AS and *α*-MGH sketches in Figure 3d. For these instances, we note that the sizes of the sketches generated by MGH and *α*-MGH observe some variability (as expected when using different hash functions). However, the variability is significantly less than that observed by AS and *α*-AS sketches. In fact, the sketches generated by *α*-AS are quite unstable, sometimes resulting in nearly empty sketches, and sometimes resulting in very large sketches compared to the median. Our counterpart *α*-MGH, on the other hand, generates sketches whose sizes are very well concentrated around the median. These results reveal that despite MGH and *α*-MGH generating slightly larger sketches (although the asymptotic growth is the same), the sketches generated by our algorithms are more stable than those generated by AS and *α*-AS.

The stability of our algorithm is further illustrated in 3e-h, where we show the estimated Jaccard using AS and MGH. For these experiments, we generated two random sets, each comprising 100,000 random alphanumeric strings, with a Jaccard similarity of 0.5. We used AS and MGH sketches to estimate the Jaccard similarity. In Figure 3e and Figure 3f, we generated the sketches using the same hash function, but used 100 different random orders to process the elements (e and f are the same setting as a and b, respectively). Here, we note that since MGH sketches are always the same, they always estimate the same Jaccard. On the other hand, when we run AS, the resulting sketches are different each time due to sensitivity to the order in which elements are processed. As a result, AS sketches estimate a range of Jaccard values. As *α*-AS sketches are more unstable, we observe that the resulting Jaccard estimate varies from 0.48 to 0.56 – effectively rendering *α*-AS non-usable to estimate similarity.

And finally, when we keep the order of processing the same but use 100 different hash functions to estimate the Jaccard of the same two sets, we find that AS and MGH have the same degree of variability in estimated Jaccard, although the *α*-variant of AS estimates Jaccard scores across a very wide range.

Together, these results highlight two reasons to prefer MGH over AS: similarity/distance scores estimated using MGH are more stable than those estimated using AS, and results prepared using MGH can be safely recreated owing to the “order invariant” property which is missing in AS.

### 6.3 MaxGeomHash samples can estimate similarity across the entire range

We next present experiments investigating the quality of estimated similarity scores using MGH and *α*-MGH samples and the steps outlined in Section 5. We show only Jaccard estimation results; experiments with other similarity scores were performed, but omitted for brevity.

We randomly generated 5,000 pairs of sets such that the Jaccard similarity between each pair varied from 0.0 to 1.0. Each individual set contains 100,000 alphanumeric strings of length 10. For each pair, we estimated the Jaccard similarity using MGH samples with parameters *b* ∈ { 70, 80, 90, 100 }, and *α*-MGH samples with parameters *α* ∈ { 0.4, 0.45, 0.5 }. These parameters were selected arbitrarily to illustrate MGH and *α*-MGH’s ability to provide for a balance between MinHash and FracMinHash. In practice, the users select the parameter to fit their need for accuracy and available computational resources. In Figure 4, we show these estimated Jaccard values against the true Jaccard scores. Estimated values using MGH samples is shown in Figure 4(a) and *α*-MGH shown in Figure 4(b). For visual clarity, only a random 1000 points are plotted.

**Fig. 4:**
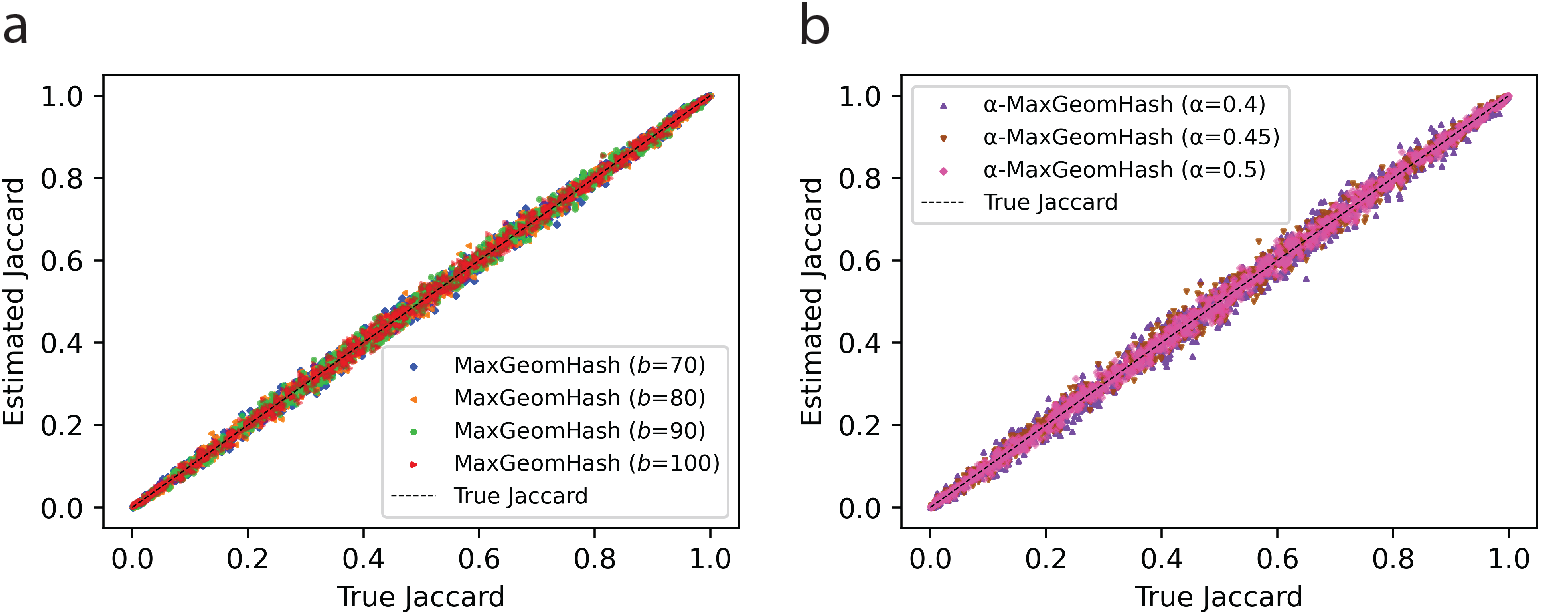
Estimated Jaccard scores against the true Jaccard scores. Estimated Jaccard scores are computed using MGH (shown in Panel a) and *α*-MGH (shown in Panel b) samples.

Estimated scores highly correlate with the true values (minimum *R*^2^ of 0.9932 over all 7 parameter settings). We note slight variability in the estimated Jaccard values, with the variance decreasing as the sketches become larger as *b* and *α* become larger. This behavior is expected in sampling/sketching techniques that trade efficiency for tolerable error. Nonetheless, estimated values remain tightly aligned with true scores without systematic bias, indicating that MGH and *α*-MGH samples provide accurate, unbiased similarity estimation across the entire range.

### 6.4 MaxGeomHash allows asymptotically unbiased similarity estimation while providing a balance between MinHash and FracMinHash

We next compare MaxGeomHash with two other widely used sketching methods, MinHash and FracMin-Hash. We generated pairs of random sets (*A, B*) where the number of elements in *A* and *B* is the same and the Jaccard similarity between *A* and *B* is fixed at 0.5. We varied the number of elements in *A* and *B* from 100K to 50M (million). We focus on slightly larger sets (in contrast to the results presented in Figure 2) to capture the range where FracMinHash sketches start to become larger than MinHash sketches.

For each of these (*A, B*) pairs, we computed sketches using four sketching algorithms: MinHash, FracMin-Hash, MGH, and *α*-MGH. For MinHash and FracMinHash, we used the default parameters (*k* = 1000, *s* = 0.001, respectively) set in the software tools Mash [22] and sourmash [9], respectively. These default parameters are widely used in many bioinformatics analyses. To compute MGH and *α*-MGH sketches, we used *b* = 90, and *α* = 0.45, respectively – which provide a good balance between MinHash and FracMinHash. We note that this specific choice of *b* = 90 and *α* = 0.45 was determined by MinHash and FracMinHash parameters. This parameter choice does not restrict the generality of the results: for any choice of parameters for MinHash and FracMinHash, we can identify *b* and *α* values for the two versions of MaxGeomHash that can offer an accuracy-efficiency balance between MinHash and FracMinHash.

For each algorithm, we ran 500 independent trials. In each trial, we computed sketches of the sets *A* and *B* using a distinct seed and estimated the Jaccard similarity using these sketches. The average size of the sketches is shown against the number of elements in *A* and *B* in Figure 5a. From the estimated values, we also computed the mean squared error, MSE (note that the true Jaccard is fixed at 0.5), and show how the MSE changes as the sets grow large in Figure 5b.

**Fig. 5:**
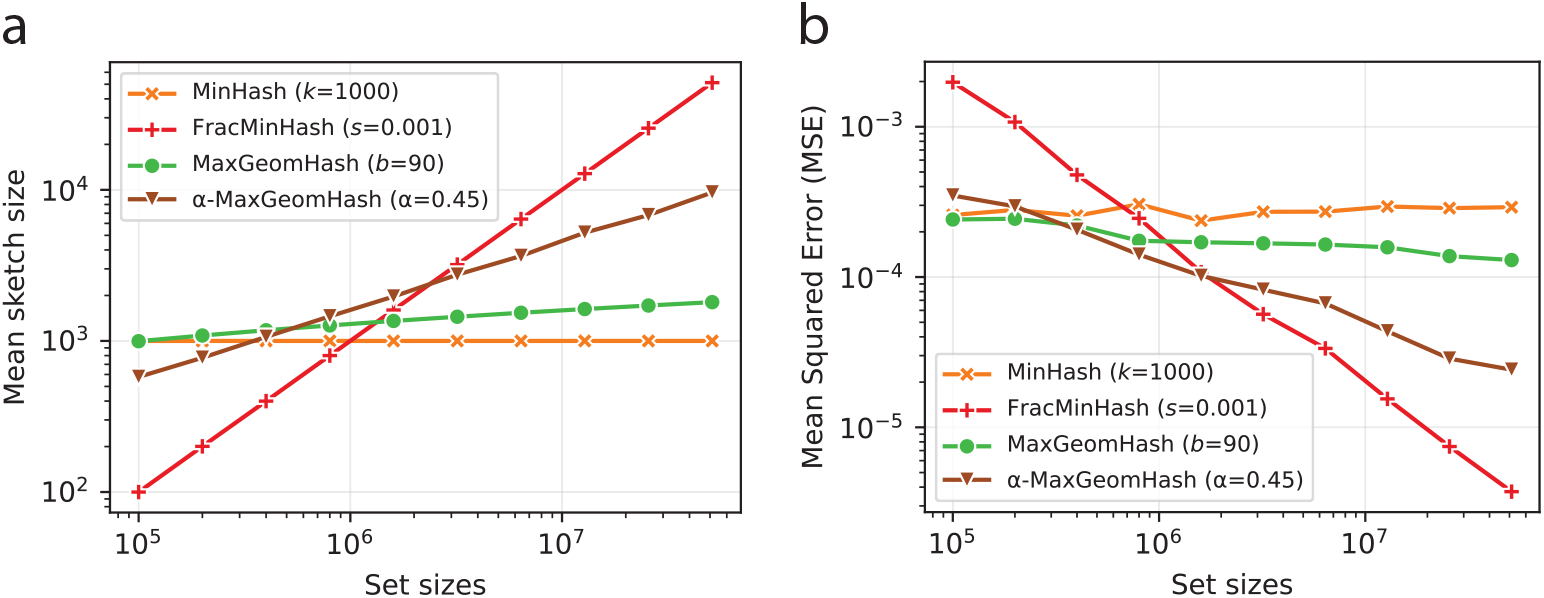
(a) Average sketch sizes for four sketching algorithms against sizes of the original sets, computed from 50 independent trials. (b) Mean squared error (MSE) in estimating the Jaccard similarity of two sets, where the sizes of the two sets are equal. The number of elements in the original sets is shown on the horizontal axis. MSE was averaged from 50 independent hash seeds.

Panels in Figure 5 show complementary results: as sketch size grows, MSE decreases. MinHash sketches are fixed in size, so MSE remains relatively constant. For the other three algorithms, sketch size grows with | *A* | and | *B* |, and MSE diminishes towards zero – indicating that FracMinHash, MGH, and *α*-MGH are asymptotically unbiased. With the fastest growing sketch size, FracMinHash exhibits the quickest decrease in MSE. MGH and *α*-MGH sketches grow slower than FracMinHash sketches, so their MSE decays less quickly, while consuming fewer computational resources than FracMinHash.

We repeated this setup for other similarity metrics (cosine, Kulczynski 2) and fixed similarity values (0.1 to 0.9, increments of 0.1), observing similar trends, but omitting for brevity.

### 6.5 An application using real biological data

We conclude with a real biological data application: estimating a similarity tree (a proxy for phylogenetic tree) among ten mammals. The phylogenetic relationships between mammals are well-established [30]. We assess whether *k*-mer-based sketching techniques can approximate the known phylogeny.

We downloaded ten mammal genome assemblies from Ensembl [32], extracted canonical *k*-mers (*k* = 31), and computed *k*-mer-sketches using four algorithms: MinHash, FracMinHash, MaxGeomHash, and *α*-MaxGeomHash. We used the same parameters as in Figure 5, with motivation detailed in Section 6.4. We computed pairwise Jaccard similarity by loading all sketches in memory, then converted Jaccard scores to genome-wide average mutation rates (i.e., one minus average nucleotide identity, ANI) assuming a simple mutation model using methods from [11]. We then computed a similarity tree from these pairwise mutation rates using UPGMA (Unweighted Pair Group Method with Arithmetic Mean), a standard distance-based method for constructing rooted phylogenetic trees under a molecular clock assumption [6, 20]. We also experimented with the Neighbor Joining algorithm and observed similar trends, but opted to present the UPGMA-generated trees for clarity. Estimated trees for each sketching method are shown in Figure 6.

**Fig. 6:**
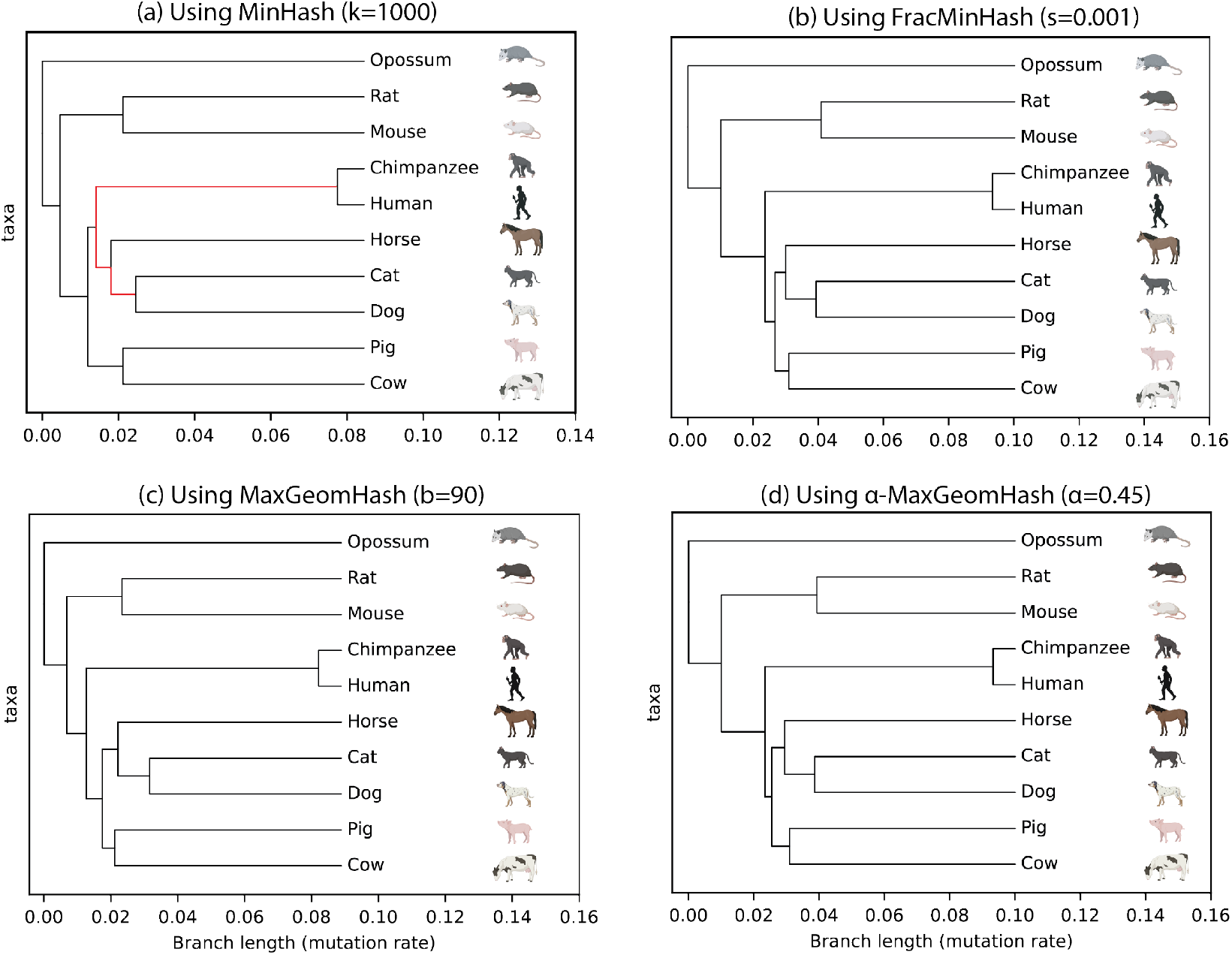
Similarity tree of ten mammal genomes estimated using four sketching methods using *k*=31-mersketches. The red branch in (a) shows that by using MinHash, Carnivora mammals Cat and Dog are placed close to Primates, whereas in reality, they resemble Pig, Cow, and Horse more, and belong in the Laurasiatheria clade. The other three methods consume more resources and correct this mistake, where MGH and *α*-MGH offer better resource usage compared to FracMinHash (time & memory listed in Table 1). Icons of the mammals are collected from BioRender.com.

All methods correctly identify Opossum as the outgroup (most distantly related to the others) and correctly group similar organisms: Rat and Mouse (Rodentia), Human and Chimpanzee (Primates), Cat and Dog (Carnivores), and Pig and Cow (Cetartiodactyla). However, all methods incorrectly place rodents and Primates far apart, though these clades are sister groups.

The major discrepancy between the MinHash tree and the others (Figure 6a) is that with only 1000 hashes, Carnivora (Cat and Dog) are placed close to Primates (Human and Chimpanzee), whereas they actually belong in Laurasiatheria with Pig, Cow, and Horse. Using larger sketches, FracMinHash, MGH, and *α*-MGH correct this error and are thus more accurate than MinHash. However, FracMinHash requires more computation due to linear sketch growth, whereas MGH and *α*-MGH reduce computational costs by producing smaller sketches that grow sub-linearly.

Computational resources are reported in Table 1. For fair comparison, we wrote a simple C++ program to read sequences from genome files, split them into *k*-mers, compute sketches, and store them as plain text files of hash values. FracMinHash is the fastest to compute despite a larger size, since its logic is simple – whether to keep a hash value doesn’t depend on other hash values. The other three methods require maintaining a min (max) heap-like data structure, resulting in slightly higher CPU time. Memory usage for computing sketches is similar across methods, with FracMinHash consuming slightly more due to larger sketches. Disk space for storing sketches starkly highlights size differences: MinHash requires the least space, FracMinHash the most. Finally, computational resources for loading sketches and computing pairwise Jaccard values (and genome-wide average mutation rates) follow sketch sizes, with MinHash consuming the least resources, FracMinHash the most, and MGH and *α*-MGH providing balance.

We conclude by highlighting that MGH and *α*-MGH, using parameters *b* = 90 and *α* = 0.45, respectively, can generate clusters that are as accurate as FracMinHash, yet they consume many times less resources when performing pairwise similarity estimation (516x and 22x faster, respectively; 167x and 22x lightweight in memory, respectively; 419x and 22x less storage, respectively). Overall, MaxGeomHash sketches can estimate a similarity tree more efficiently than FracMinHash, and more accurately than MinHash when the parameters are selected carefully. We note that while we used similarity (phylogenetic) tree estimation as an example to highlight this advantage of MaxGeomHash, such benefits can be had in any analysis where sketches are used to estimate similarity/distance.

## 7 Conclusion

We introduced MaxGeomHash and *α*-MaxGeomHash, both sketching algorithms are one-pass, dependable, order-independent, and mergeable. Without knowing the number of distinct elements *n*, MGH yields an expected distinct element sample size of *Θ*(*b* log(*n/b*)) while *α*-MGH yields *Θ*(*n*^*α*^). This situates MGH and *α*-MGH directly between MinHash (constant sketch size) and FracMinHash (linear sketch size), obtaining the more accurate scaling feature of FracMinHash with the advantage of smaller sketch sizes. We analyzed sample-size expectation/variance and obtained (asymptotically) unbiased estimators for the Jaccard index, and leveraged recent theory results that automatically extend these estimators to a variety of other metrics (containment, cosine, Kulczynski 1, etc.) as well as provide explicit error bounds and concentration inequalities. These properties make MGH and *α*-MGH well-suited for the diverse sketching applications that form a core substrate for sequence search, clustering, quality assessment, phylogenetics, and metagenomic surveillance. Workflows that currently use FracMinHash or MinHash (such as Mash screen [21], sourmash gather [12], Skani [27], YACHT [15], fmh-funcprofiler [10], sylph [28], etc.) could be retooled to use MGH to shrink memory/IO budgets without sacrificing accuracy guarantees.

Similarly, recent work relating sketching based metrics to biologically informative metrics like ANI [11], AAI [23], and dN/dS [24] translate directly to MGH: one merely replaces a FracMinHash estimate of these metrics with the MGH counterpart. Recent large-scale computational biology projects, like the Logan effort [2] prioritize storage efficiency. Utilizing MGH in such infrastructures and projects can substantially reduce persistent index size with only modest (and exactly quantifiable) accuracy changes, while retaining compatibility with downstream operations on such sketches.

Future improvements include refining our C++ implementation which is not yet fully optimized. There remains headroom from cache-aware bucket layouts, succinct per-bucket priority structures, SIMD-friendly hash partitioning, and task-parallel refinement. These engineering improvements should further reduce run-time and memory while preserving the theoretical guarantees established for MGH and *α*-MGH.

## Acknowledgments

The research of David Koslicki has been supported by NIH NIGMS project R01GM146462. The research of Conrado Martínez has been supported by funds from the MOTION Project (Project PID2020-112581GB-C21) of the Spanish *Ministerio de Ciencia e Innovación* (MCIN/AEI/10.13039/501100011033), and the PLASMA Project (Project ANR-25-CE48-3555) of the French *Agence Nationale de la Recherche*. The work was initiated when Mahmudur Rahman Hera was a PhD student at Penn State. During this time, Mahmudur Rahman Hera was supported by NIH NIGMS project R01GM146462. The authors gratefully acknowledge Dr. Anat Kreimer for supporting the completion of this work during Mahmudur Rahman Hera’s postdoctorate appointment at Rutgers.

## A Analysis of 𝔼 [1/*S*_*n*_]

The variance of the similarity estimators discussed in Section 5 is in all cases *Θ*(𝔼 [1*/S*_*n*_]), where we write *S*_*n*_ for the size of the samples, stressing the fact that the random variable depends on *n*. Moreover, for similarity measures such as cosine or Kulczynski 1, we have asymptotically unbiased estimation, with the bias of order *Θ*(𝔼 [1*/S*_*n*_]).

It is therefore important to know the behavior of 𝔼 [1*/S*_*n*_] as *n*→ ∞ ; we show that for the two variants of MGH in this paper we have

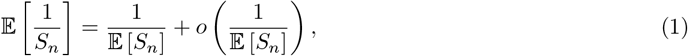

and since 𝔼 [*S*_*n*_] → ∞ as *n* → ∞, we conclude that the variance tends to 0 for large *n*, and likewise the bias, if any, also tends to 0.

To show (1) we will use the following: 1) *S*_*n*_ is a positive integer value, 0 *< S*_*n*_ ≤ *n*, whenever *n >* 0; 2) *µ*_*n*_ = 𝔼 [*S*_*n*_] is a non-decreasing function of *n* and tends to + ∞ when *n*→ ∞ ; 3) 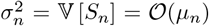 for both variants of MGS: it is 𝕍 [*S*_*n*_] = *Θ*(*b*) for standard MGH, and 𝕍 [*S*_*n*_] = *Θ*(*n*^*α*^) for *α*-MGH.

Applying Chevbyshev’s inequality, we immediately get that *S*_*n*_ has concentration around its mean—we write *S*_*n*_ = *µ*_*n*_ + 𝒪_P_(*σ*_*n*_), that is, for any *ϵ >* 0, there exists *δ* := *δ*(*ϵ*) such that

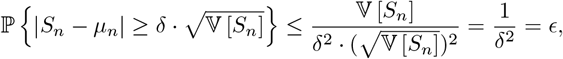

using 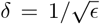. Because of (2) and (3) above, we have that 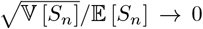 as *n* → ∞ and *S*_*n*_ = *µ*_*n*_ + 𝒪_P_(*σ*_*n*_) are equivalent.

Define now *X*_*n*_ := *S*_*n*_ − *µ*_*n*_, so that 𝔼 [*X*_*n*_] = 0 and 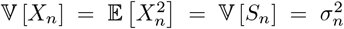. Note that −*n* ≤ *X*_*n*_ ≤ *n* for all *n >* 0. Using a Taylor expansion of 1*/S*_*n*_ around *µ*_*n*_:

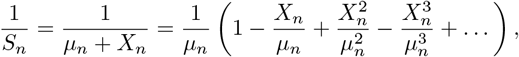

and taking expectations, we obtain

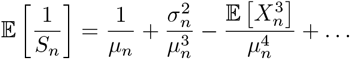

The second-order term 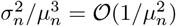 and higher-order terms are negligible, because the concentration *S*_*n*_ = *µ*_*n*_ + 𝒪_P_(*σ*_*n*_) and −*n* ≤ *X*_*n*_ ≤ *n* guarantee 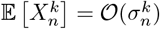. for all *k* ≥ 2.

## B Methods

In this section, we give the implementation details that were not included in the main text for the sake of brevity.

### B.1 Implementation of MaxGeomHash and *α*-MaxGeomHash

We implemented MaxGeomHash and and *α*-MaxGeomHash both using Python and using C++. The Python implementation was used for all the simulation experiments (experiments presented in Sections 6.1 to 6.4). The C++ implementation was used to compute the *k*-mer-sketches of the mammal genomes (experiments presented in Section 6.4). In the Python implementation, we used MurMurHash3 function implemented in the Python package “mmh3” [26]. We used 64-bit unsigned hash values for our purposes, which is a common practice in bioinformatics applications [4,9]. To maintain the items with the *b*-largest hash values in the non-empty buckets, we used the package “heapq” in Python. For our C++ implementation, we used a public C++ implementation of the same hash function, and also used unsigned 64-bit hash values. We used priority queues, implemented in the standard template library of C++, to maintain the non-empty buckets in our C++ implementation.

### B.2 Obtaining the mammal genomes

We obtained the ten mammal genomes from Ensembl [32] Release-112. A list of these mammals and their Ensembl IDs is as follows.

### B.3 Storing *k*-mer-sketches as plain text files

We store all sketches in a simple, human-readable plain-text format consisting of a metadata header followed by algorithm-specific data records. Each file begins with several comment lines (prefixed by ‘#’) that record the sketch version, algorithm name, *k*-mer size, hash seed, hash function, and the parameters relevant to the particular sketching algorithm. After the header, each file lists one record per line. For the MaxGeomHash and *α*-MaxGeomHash sketch, each line contains four comma-separated fields: the bucket index, the primary hash, a secondary hash used by the geometric sampling process, and a frequency count. This representation encodes all records in the sample produced by MaxGeomHash.

The bottom-*k* and FracMinHash sketches use simpler formats, each storing a single hash value per line. For bottom-*k*, the file contains the *k* smallest hash values observed in the sequence, as specified by the ‘params.k’. For FracMinHash, the file lists all hashes that fall below a threshold derived from the sampling scale parameter (‘params.scale’), effectively implementing downsampling by a fixed fraction. This minimalistic and uniform design across algorithms ensures ease of debugging and straightforward integration into similarity estimation.

### B.4 Comparing pairs of *k*-mer-sketches

When comparing two sketches, we first check if their metadata indicate compatibility – if hash function or seeds differ, or if *k*-mer sizes differ, or if the parameters of the sketching algorithm employed to compute the sketches differ, then the two sketches are not compatible with each other, and the similarity estimation does not proceed further. If the two sketches are found to be compatible, the algorithm-specific rules are used to compute a similarity metric, such as the cosine similarity or the Jaccard index.

### B.5 Recording running times, memory usages, and storage usages

We used the standard “/usr/bin/time -v” command to record the CPU time usage, as well as the peak memory usage (maximum resident set size). For the storage usage, the bash utility “find” was used to get the total space required to store the sketches in plain text files.

## Footnotes

5 Both MinHash and bottom-*k* render samples of fixed size, say *M*. MinHash actually returns *M* hash values, not the elements themselves: it applies *h*_1_, …, *h*_*M*_ hash functions to each item in the data set, and keeps the minimum values observed for each of the *M* hash functions. In bottom-*M*, we apply a hash function *h* to each item *x* in the data set, and we keep the *M* smallest observed hash values. Despite the two algorithms are different (specially in terms of practical efficiency) they render equivalent random samples of size *M*, and we will loosely use the terms MinHash and bottom-*k* interchangeably.

6 Assuming repetitions are filtered out.

7 It holds for any variant of the algorithm, whether the capacity of the buckets is *b*, ⌈2^*βi*^⌉ or any function *f*_*i*_.

